# From maternal glucocorticoid and thyroid hormones to epigenetic regulation of gene expression: an experimental study in a wild bird species

**DOI:** 10.1101/2023.03.07.531470

**Authors:** Mikaela Hukkanen, Bin-Yan Hsu, Nina Cossin-Sevrin, Mélanie Crombecque, Axelle Delaunay, Lotta Hollmen, Riina Kaukonen, Mikko Konki, Riikka Lund, Coline Marciau, Antoine Stier, Suvi Ruuskanen

## Abstract

Offspring phenotype at birth is determined by its genotype and the prenatal environment including exposure to maternal hormones. Variation in both maternal glucocorticoids and thyroid hormones can affect offspring phenotype. However, the underlying molecular mechanisms shaping the offspring phenotype, especially those contributing to long-lasting effects, remain unclear. Epigenetic changes (such as DNA methylation) have been postulated as mediators of long-lasting effects of early-life environment. In this study, we determined the effects of elevated prenatal glucocorticoid and thyroid hormones on handling stress response (breath rate), DNA methylation and gene expression of glucocorticoid receptor (GCR) and thyroid hormone receptor (THR) in great tit (*Parus major*). Eggs were injected before incubation onset with corticosterone (main avian glucocorticoid) and/or thyroid hormones (thyroxine and triiodothyronine) to simulate variation in maternal hormone deposition. Breath rate during handling and gene expression of GCR and THR were evaluated 14 days after hatching. Methylation status of GCR and THR genes were analyzed from the longitudinal blood samples taken 7 and 14 days after hatching, as well as in the following autumn. Elevated prenatal corticosterone level significantly increased the breath rate during handling, indicating enhanced stress response and/or metabolism. Prenatal corticosterone manipulation had CpG-site-specific effects on DNA methylation at the GCR putative promoter region, while it did not significantly affect GCR gene expression. GCR expression was negatively associated with earlier hatching date and chick size. THR methylation or expression did not exhibit any significant relationship with the hormonal treatments or the examined covariates, suggesting that TH signaling may be more robust due to its crucial role in development. This study supports the view that maternal corticosterone may influence offspring metabolism and stress response via epigenetic alterations, yet their possible adaptive role in optimizing offspring phenotype to the prevailing conditions, context-dependency, and the underlying molecular interplay needs further research.

## 1. Introduction

Maternal effects occur when parental phenotype directly affects the offspring phenotype (Bernardo 1996). These effects may persist throughout one’s lifetime and even to subsequent generations (Bernardo 1996; Mousseau & Fox 1998). Maternal effects may have adaptive value when mothers’ experience of the environment is transmitted to the next generation (for example via molecular markers, hormones, resources or care), and when this increases offspring fitness (Mousseau & Fox 1998; Love & Williams 2008; Weber et al. 2018; Yin et al. 2019; Zhang et al. 2020). However, maternal hormonal effects may also be due to mere physiological constraints, and their adaptive role remains debated (Groothuis et al. 2005; Sánchez-Tójar et al. 2020; Zhang et al. 2020). Hormones are one of the main mechanisms of maternal effects because they affect various traits by altering gene expression and cellular functions (Groothuis et al. 2005; Podmokla et al. 2018; Groothuis et al. 2019). Since environmental factors such as food abundance alter maternal hormone production and transport to the offspring, maternal hormones may program the offspring to better cope with the prevailing environmental conditions (Groothuis et al. 2019).

Maternal hormonal effects have been widely explored in oviparous species, such as birds. Since the embryo develops outside of the mother’s body, embryonic hormones can be measured and manipulated uncoupled from the mother’s physiology (Groothuis et al. 2019). Maternally derived steroid hormones have been found to have long-lasting or even transgenerational effects on postnatal phenotype and fitness-related traits such as growth and reproduction (Groothuis et al. 2005; Podmokla 2018). In birds, both glucocorticoids and thyroid hormones are transferred from the mother’s blood to the egg yolk, leading to both transient and/or long-lasting phenotypic changes in the offspring (Hayward & Wingfield 2004; Schoech et al. 2011; Ruuskanen & Hsu 2018). For instance, great tits (*Parus major*) hatching from eggs with experimentally elevated corticosterone (main avian glucocorticoid) levels exhibit prolonged begging behavior and increased breath rates (Tilgar et al. 2016). Breath rate is at least partly controlled by the autonomic nervous system and reflects indirectly metabolism, but also stress response (Carere & van Oers 2004; Careau *et al*. 2008; Yackle *et al*. 2017), and it shows repeatability, heritability, and significant variation between individuals at handling (Fucikova et al. 2008). The effects of *in ovo* corticosterone manipulations have been found to alter the hypothalamus-pituitary-adrenal (HPA) axis responses, which is crucially involved in stress physiology (Love and Williams 2008; Haussmann et al. 2012; Tilgar et al. 2016). Differences in the behavior, metabolism, and stress response may represent different strategies to cope with environmental challenges (Koolhaas et al. 1999; Carere et al. 2001; Romero 2004). As the most advantageous coping strategy can be dependent on the prevailing environmental conditions (*e.g.* food abundance, predation pressure, population density, weather unpredictability; Koolhaas et al. 1999; Carere et al. 2005; Careau et al. 2008), prenatal exposure to maternal hormones may be important in preparing the offspring for the expected environmental conditions after hatching to maximize fitness prospects. Yet — regardless of whether maternal hormonal effects are adaptive or not, the molecular mechanisms underlying effects on phenotype remain poorly understood (Groothuis & Schwabl 2008; Groothuis et al. 2019; Bentz 2021).

Changes in the epigenetic regulation (*i.e.* mechanisms translating the information of a genotype to various phenotypes; Waddington 1942) of the hormonal pathways could be one such mechanism (Jimeno et al. 2019; Ruiz-Raya et al. 2023). One of the best-studied epigenetic mechanisms is DNA methylation, which facilitates changes in gene expression, imprinting, and transposon silencing (Jaenisch & Bird 2003, Vogt 2021, Laine et al. 2022), and is known to be affected by age, environmental quality, and physiological condition (De Paoli-Iseppi et al. 2018; Lindner et al. 2021; Mäkinen et al. 2022). Epigenetic alterations may function as a tool for individuals to acclimatize to changing environmental conditions, but they may also mediate transgenerational adaptation (Guerrero-Bosagna et al. 2018; Sepers et al. 2021; Vogt 2021). Epigenetic alterations of the HPA-axis, especially hormone receptor genes such as glucocorticoid receptor (GCR, transcribed by *NR3C1)*, have been suggested to mediate the effects of prenatal exposure to glucocorticoids (Groothuis & Schwabl 2008; Ahmed et al. 2014; Zimmer & Spencer 2014). Indeed, high concentrations of *in ovo* corticosterone have been found to increase offspring GCR methylation and decrease the receptor protein expression in chicken (*Gallus gallus domesticus*) hypothalamus (Ahmed *et al*. 2014). The role of receptor methylation in mediating the effects of prenatal thyroid hormones has received less attention in previous studies: Van Herck *et al*. (2012) discovered that TH supplementation to chicken egg yolk increased TH concentration in the brains of chicken embryos 24h after manipulation and altered TH membrane transporter expression; however, the effects on thyroid hormone receptor (THR, transcribed by *THRA* and *THRB*) expression or methylation have not been investigated. The few studies (Ahmed et al. 2014; Zimmer & Spencer 2014; Van Herck *et al*. 2012) that have addressed the epigenetics of prenatal hormonal effects by directly manipulating the egg hormone concentrations have (1) mainly focused on glucocorticoids, and (2) hormone manipulation occurred when incubation already started. Manipulations during incubation do not mimic maternal deposition and could lead to different effects since CORT and TH are likely to be at least partly metabolized by the growing embryos (CORT: Vassallo et al. 2019; TH: Ruuskanen et al. 2022). Additionally, the impact on methylation pattern has only been examined cross-sectionally for a relatively short period after hatching so far, which precludes to evaluate the potential long-lasting nature of epigenetic changes induced by maternal hormonal effects, knowing that human studies have shown both consistency and flexibility in methylation patterns with time within an individual (Komaki et al. 2021).

To fill these gaps, we experimentally elevated glucocorticoid (i.e. corticosterone) and thyroid (i.e. triiodothyronine,T3 and thyroxine, T4) hormone concentrations within the natural range in wild great tit (*Parus major*) eggs before incubation onset to simulate the causal effects of maternally elevated hormones on offspring phenotype. Breath rate in response to handling was calculated as an indirect measure of acute stress response and metabolism (Carere et al. 2001; Carere & van Oers 2004; Fucikova et al. 2008). We then investigated possible alterations in the methylation status of the glucocorticoid and thyroid hormone receptors using a longitudinal study design (blood sampling of the same individuals at days 7 and 14 post-hatching, as well as early adulthood), and assessed gene expression patterns at a single time-point (day 14 post-hatch).

Considering their effect on stress response and metabolism, we predict that the prenatal glucocorticoid and thyroid hormone exposure will increase the offspring breath rate during handling, (*reviewed in* Thayer et al. 2018). Given that hormonal responses are regulated by a negative feedback system (McNabb 2007; Cottrell & Seckl 2009), higher prenatal exposure and possibly enhanced intrinsic hormonal production during early development may have increased the receptor gene methylation and downregulated its mRNA expression. These predictions are supported by an association between high baseline plasma corticosterone and low GCR expression in zebra finches (*Taeniopygia guttata,* Jimeno et al. 2019), as well as by the fact that prenatal stress has been found mainly to decrease GCR expression (*reviewed in* Kapoor et al. 2006; Cottrell & Seckl 2009, yet see Zimmer et al. 2017). We predict decreased methylation patterns with age, following Gryzinska et al. (2013), who reported decreased global methylation in chicken blood between one-day-old chicks and 32-week-old hens (yet see De Paoli-Iseppi et al. 2018). However, directional predictions of the age-related methylation changes in our target genes need to be approached with caution; all genes may not follow the global pattern and could even exhibit opposite patterns to global trends (De Paoli-Iseppi et al. 2018).

## 2. Materials and methods

### 2.1. Study population

The study was conducted in a wild great tit nest box population (374 boxes in 10 plots, all within 3 km range) in Ruissalo, Southwestern Finland (60° 26’ N, 22° 10’ E). Nesting activity was monitored every four-five days. The nests where egg-laying had started were visited daily until hormone manipulation was completed. The experimental protocol has been described in detail in (Cossin-Sevrin et al. 2022): this study concerns a subsample of the nests included in the larger study.

### 2.2 Hormone manipulations

The mean final clutch size of great tit nests was 8.2 (SD=1.83), ranging from 2 to 10. All eggs from one nest received the same treatment. The nests were assigned to four treatment groups: control (CO, N=11 nests), glucocorticoid hormone supplementation (CORT, N=12), thyroid hormone supplementation (TH, N=12), and a combination of glucocorticoid and thyroid hormone (CORT+TH; N=10). The treatments were assigned to the nests randomly, but sequentially so that all treatments would be equally distributed across the breeding season, and attention was given to geographical distribution (i.e., all treatments present in all forest plots). Egg injection started on the day of the 5^th^ egg (as females may start to incubate before clutch completion) and every day thereafter for newly-laid eggs, which ensured that no incubation had occurred at the time of the injection. The doses of TH and CORT were chosen to increase the average content in the yolks by 2SD (Podmokla et al. 2018). Each egg in the TH group was injected with a combination of 0.32 ng of T4 and 0.04 ng of T3. For the corticosterone treatment group, 0.2 ng was injected per egg. The combination group (CORT+TH) received the sum of all hormones (0.32 ng T4, 0.04 ng T3 and 0.2 ng corticosterone). Each egg received an injection of 2 µl, containing the hormone in question, dissolved in 0.1 mol/l NaOH (TH) or 99% ethanol (CORT), and diluted in 0.9% NaCl. For more details on the injection procedure, see Cossin-Sevrin *et al*. (2022).

### 2.3 Phenotypic measurements

The nests were visited daily starting 2 days before the predicted hatching to record the hatch date (= day 0). Nestlings were visited 2, 7, and 14 days after hatching for identification, phenotypic measurements (weight, wing length) and blood sampling (day 7 and 14) as described in Cossin-Sevrin *et al*. (2022). Additionally, 14 days after hatching, we measured handling stress response as a proxy of different stress-related coping strategies (Carere & van Oers 2004; Fucikova et al. 2008) by calculating the individual’s breath rate (see below) in response to handling (Carere & van Oers 2004; Fucikova *et al*. 2008; Liang *et al*. 2018). Breath rate was measured from one to three randomly-selected chicks per nest, immediately after taking each individual from the nest (before other measurements) following the protocols described in Carere and van Oers (2004) and Fucikova *et al*. (2009). The breath rate was measured as breast movements during a 60-second time. The entire measurement lasted for 75 seconds per individual (15s interval x4, 5s in between). Breath rate was calculated as the sum of breast movements during the four intervals. Sex of the individuals was determined from 14d blood samples using a qPCR approach adapted from Ellegren and Sheldon (1997) and Chang *et al*. (2008). Details are described in Cossin-Sevrin *et al*. (2022).

Individuals were recaptured as juveniles (ca. 9–20 weeks after fledging) in the following autumn: 20 of the 39 individuals included in the methylation analysis were recaptured. Mist nets (with playback) were set up in 7 feeding stations across the study area. Each feeding station was visited for 3 hours per netting on 3 distinct days during October-November (total of 100 hours of mistnetting). Weight and wing length measurements, as well as blood samples were collected for juvenile individuals applying the same workflow as above.

### 2.1 Methylation analysis

#### 2.1.1 DNA methylation: DNA extraction

DNA methylation of glucocorticoid receptor gene *NR3C1* and thyroid hormone receptor B *(THRB)* were detected by bisulfite conversion followed by pyrosequencing. For each nest, one randomly selected individual was used in the methylation analysis. DNA was extracted from the frozen blood samples of 40 great tit individuals, each of which were sampled longitudinally 2 or 3 times (Table 2). The DNA samples were stored at −80 °C after extraction. DNA extraction and quality assessment are described in Cossin-Sevrin et al. (2022).

**Table 1.** Number of individuals and nests by treatment group in the whole experiment and for different analyses (breath rate, methylation, gene expression). Numbers of nests are given in brackets. Treatment groups are coded as follows: CO=control; CORT=corticosterone; CORT+TH=corticosterone and thyroid hormone combination group; TH=thyroid hormone. There were 1-3 randomly selected individuals per nest for the breath rate analysis (36 nests). One randomly selected individual per nest was included in the methylation and gene expression analyses. All individuals in the gene expression analyses were included in the methylation analyses. We tried to also maximise the overlap between data on breath rate, methylation and gene expression from the same individuals, but this was not always possible due to limited blood sample availability. In the end, 18 individuals with methylation data had also breath data data, and 16 individuals with gene expression data had also breath rate data. There were 45 different nests in total (CO=11; CORT=12; CORT+TH=10;TH=12).

|  | *In ovo treatment* |  |  |  |  |
| --- | --- | --- | --- | --- | --- |
|  | CO | CORT | CORT+TH | TH | Total |
| N breath rate | 27 (10) | 20 (10) | 16 (7) | 20 (9) | 83 |
| N methylation | 10 (10) | 10 (10) | 9 (9) | 10 (10) | 39 (39) |
| N gene expression (*NR3C1/THRA*) | 10/10 (10) | 8/9 (9) | 6/6 (6) | 9/8 (9) | 34 (45) |

**Table 2.** Sample sizes as number of individuals/CpG-sites included in the methylation analysis after quality filtering for each gene (NR3C1=glucocorticoid receptor; THRB=thyroid hormone receptor β), treatment group (CO=control; CORT=corticosterone; CORT + TH=corticosterone and thyroid hormone; TH=thyroid hormone) and age (DAH=days after hatching).

| NR3C1 |  |  |  |  |  |
| --- | --- | --- | --- | --- | --- |
| Age | CO | CORT | CORT + TH | TH | Total |
| 7 DAH | 10/119 | 9/108 | 9/106 | 10/112 | 38/445 |
| 14 DAH | 10/112 | 10/120 | 8/95 | 10/117 | 38/444 |
| Juvenile | 5/59 | 5/60 | 5/54 | 5/60 | 20/233 |
| Total | 25/290 | 24/288 | 22/255 | 25/289 | 96/1122 |
| THRB |  |  |  |  |  |
| Age | CO | CORT | CORT + TH | TH | Total |
| 7 DAH | 10/40 | 10/39 | 8/32 | 10/39 | 38/150 |
| 14 DAH | 10/39 | 10/40 | 7/28 | 10/40 | 37/147 |
| Juvenile | 5/18 | 4/16 | 5/20 | 5/20 | 19/74 |
| Total | 25/97 | 24/95 | 22/80 | 25/99 | 96/379 |

#### 2.1.2 DNA methylation: Bisulfite conversion

Bisulfite conversion of the DNA samples was conducted by using EpiTect Fast DNA Bisulfite Kit (Qiagen, cat. 59824) and by following the manufacturer’s high concentration protocol. For each sample 20 µl of genomic DNA (10 ng/µl) was used as a starting material. The cleaned bisulfite converted DNA samples were stored at 4 °C, and the following PCR was conducted within 24 hours.

Thyroid hormone receptors are coded by two genes: alfa and beta. The epigenetic regulation of *THRB* has previously been shown to be involved in many human phenotypes such as cancer, obesity, and aging (Joseph et al. 2007; Ling et al. 2010; Kim et al. 2013; Pawlik-Pachucka et al. 2018; Shimi et al. 2022), and affected by exposures to e.g. thyroid hormones and environmental toxins in mice (Cho et al. 2021; Laufer et al. 2022), and thus we were interested in investigating if that is the case also in the context of avian maternal hormones, and chose *THRB* for the methylation analyses.

Chromosomes 13 (for *NR3C1*, GenBank assembly accession GCA_001522545.3) and 2 (for *THRB*, GenBank assembly accession CM003710.1) of the great tit genome (GenBank assembly accession GCA_001522545.3) were retrieved from NCBI’s repository (NCBI Resource Coordinators 2016; Yates *et al*. 2019). For both genes, *NR3C1* and *THRB* (transcript variant X5), a region from 1800 base pairs upstream to 100 base pairs downstream of the transcription start site was selected as the putative regulatory region as with zebra finches in a study by Jimeno *et al*. (2019) on *NR3C1*. Within this region, primers were designed to amplify CpG-dinucleotide dense regions with PyroMark Assay Design Software 2.0. Primers were validated in PCR with bisulfite-treated samples (8 samples tested) and gel electrophoresis (1.5%, 90 V). Primer characters are shown in Table 3.

**Table 3.**
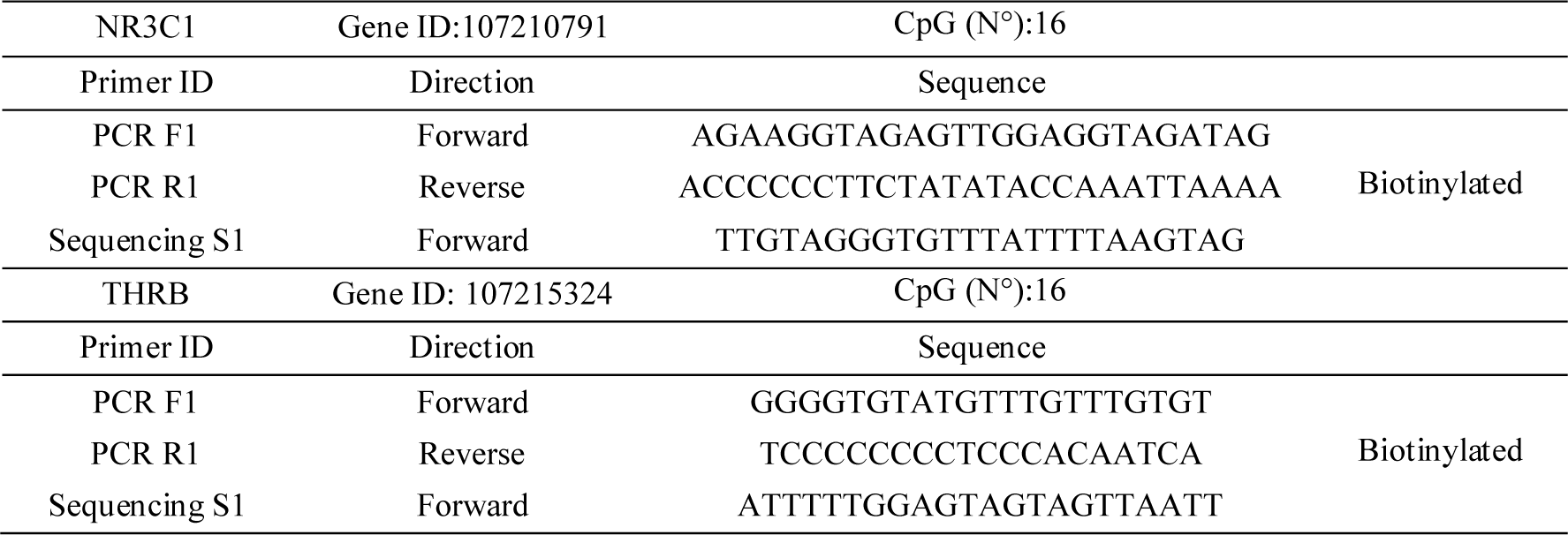
. Primers sequences used to detect the methylation status of the putative promoter regions of the glucocorticoid receptor gene (*NR3C1*) and thyroid hormone receptor gene β (*THRΒ*). Forward and reverse primer sequences are presented with the number of CpG-sites (N°) within each sequence to analyze.

#### 2.1.3 DNA methylation: PCR

The target regions of the genes of interest were prepared for amplification by using PyroMark PCR Kit (Qiagen, cat. 978903) following the manufacturer’s protocol. Bisulfite-treated DNA (4 μl, 10ng/μl) was added to the reaction mixture (without optional MgCl_2_). The thermal cycler (Applied Biosystems 2720) program varied according to the target gene: 15 min at 95°C, 45 cycles of 20 s at 94°C, 30 s at 56°C (*NR3C1*) or 60°C (*THRB*), 30 s at 95°C, and a final extension of 10 min at 72°C. The concentration (>20ng/μl) of the amplified samples and the negative controls were measured with NanoDrop ND-1000, Thermo Scientific. Samples were frozen (−20 °C) after PCR, and pyrosequencing was conducted within 3 weeks.

#### 2.1.4 DNA methylation: Pyrosequencing

For pyrosequencing, (*NR3C1* and *THRΒ*), all the samples (N=99 individuals, a total of 198 samples) were analyzed in 5 batches. All samples from the same individual at different ages were in the same batch, and the treatments were distributed as evenly as possible between the batches. One pyrosequencing run included only one gene assay (*NR3C1*/*THRΒ*).

Pyrosequencing was conducted by using PyroMark Q24 Advanced CpG Reagents (Qiagen, cat. 970922) and with PyroMark Q24 Pyrosequencing instrument (Qiagen), following the manufacturer’s protocol using 15 µl of PCR product and 2 µl Streptavidin Sepharose High Performance beads (GE Healthcare, cat. GE17-5113-01).

#### 2.1.5 DNA methylation: Quality filtering

Pyrosequencing results were first assessed in PyroMark Q24 Advanced Software (3.0.1). The pyrosequencing results included the methylation percentage and quality ranking for each CpG-site (N*_NR3C1_*=16, N*_THRΒ_*=16) within each sample (N_Sample_ = 99, in total 1584 sites for both genes where methylation was detected). Sites where the quality was classified as “Failed” by the software were discarded (*NR3C1*: 205/1584 discarded; *THRΒ*: 703/1584 discarded. As the quality of the methylation percentages decreased towards the 3’ end of the sequence, most of the discarded “Failed” data was in the 3’ end of the analyzed sequence. To ensure no sites were significantly underrepresented in the analysis, all CpG-sites with data from less than 85 samples (=86%) were discarded from the analysis (*NR3C1* 4/16 CpG-sites discarded, for *THRΒ* 12/16 CpG-sites discarded). All the methylation percentage observations with clearly deviating residuals after fitting the statistical models were considered as technical outliers and thus discarded (*NR3C1*: 3/1125 observations, DNAm% range with outliers 0-24.5%, without outliers 0-5.09%, i.e. >9 SD; *THRΒ* none).

After quality filtering, there were data from 12 CpG-sites of 96 samples from 39 individuals for the glucocorticoid receptor gene *NR3C1* (Table 2). For *THRΒ*, there was data from 4 CpG-sites of 96 samples of the same 39 individuals (Table 2). Sex ratios per treatment group are given in Table S1.

### 2.2 Gene expression analysis

The expression of glucocorticoid receptor gene (*NR3C1*) and thyroid hormone receptor genes (*THRΑ* and *THRΒ*) was examined with RT-qPCR (reverse transcription quantitative PCR) following the MIQE guidelines (Bustin *et al*. 2009). *THRB* could not be properly quantified (qPCR quantification cycle values >30, which may be due to an absence of expression in blood cells); thus the final analysis included *NR3C1* and *THRA* as genes of interest and two reference genes (*SDHA, RPL13*).

#### 2.2.1 Gene expression: RNA isolation and reverse transcription

RNA was successfully isolated from blood cells of 34 14-day-old great tits with NucleoSpin RNA Plus Kit (Macherey-Nagel). Packed blood cells (10 µl per sample) were transferred to lysis buffer and homogenized with a sterile micro-pestle, after which the remaining steps were conducted following the manufacturer’s protocol with a final elution in 50 µl of RNase-free H_2_O. The purity and concentration of extracted RNA were measured with a spectrophotometer (ND-1000, Thermo Scientific). Absorbance ratios 260/280 > 1.8 and 260/230 > 1.8 were considered thresholds for purity. Samples with RNA concentration less than 25ng/µl concentration were re-extracted from the original samples when possible. RNA integrity was validated using gel electrophoresis (E-Gel 2%, Invitrogen) and the ribosomal RNA 18S vs. 28S bands intensities. Samples not fulfilling the above-mentioned quality criteria were discarded (6/40). Isolated RNA samples were stored at −80 °C for three weeks before reverse transcription. For each sample, 600 ng of isolated RNA was reverse transcribed to complementary DNA (cDNA) with SensiFAST cDNA Synthesis -kit (Bioline) following the manufacturer’s protocol. Reverse transcribed cDNA samples were stored at 4 °C and were analyzed in qPCR within a week.

#### 2.2.2 Gene expression: RT-qPCR primers

The primers for the quantitative PCR are shown in Table 4. Primers for the great tit glucocorticoid receptor (*NR3C1,* NCBI ID: 107210791) were designed by Casagrande *et al*. (2020). Thyroid hormone receptor α (*THRΑ,* NCBI ID: 107215324) primers were designed using Primer-BLAST (Ye *et al*. 2012) using the great tit genome (GenBank assembly accession GCA_001522545.3). The reference genes used in the analyses were *SDHA* (succinate dehydrogenase complex flavoprotein subunit A, NCBI ID: 107200805) and *RPL13* (ribosomal protein L13, NCBI ID: 107209800), for which the primers were designed and validated by Verhagen *et al*. (2019).

**Table 4.**
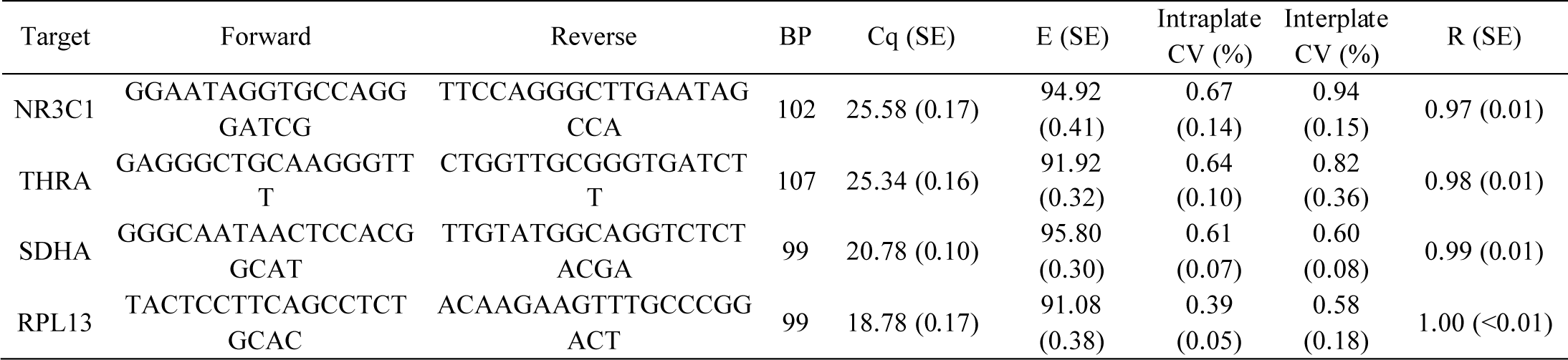
Primer sequences and performance for the genes of interest (NR3C1 and THRΑ) and reference genes (SDHA and RPL13) used in RT-qPCR. Forward and reverse primer sequences are presented with expected amplicon lengths (BP). Cq is the average quantification cycle value for each gene with associated standard error (SE). E refers to the average amplification efficiency calculated by LinRegPCR algorithm described by Ramakers et al. (2003) with the formula: *E* = 10^*slope*^ – 1. The slope is determined with linear regression from the amplification curve. Intraplate CV (%) is the average coefficient of variation for the duplicate samples, and interplate CV (%) is the coefficient of variation for two repeated qPCR plates. Technical repeatability (R) for duplicate samples is also given.

Primers for qPCR were first validated using pooled cDNA from four distinct great tit individuals that were not included in the final analysis. Primer specificity and optimal annealing temperature were confirmed by ensuring each primer produced a single narrow peak in the melt curve. NT-controls (sterile MQ-H_2_O) and template RNA (no reverse transcription) were confirmed to show no amplification before at least 5 cycles after the higher Cq of the samples of interest. A two-fold serial dilution of template cDNA from 1.5 ng to 24 ng was used to create a standard curve to evaluate primer efficiency at a wide range of starting RNA concentrations. A high-resolution melt curve analysis was used to assess the uniformity of the amplified DNA sequences as the dissociation behavior of the double stranded DNA depends on the DNA sequence. Gel electrophoresis was used to ensure a single PCR product of the expected length for a random subset of samples. Reference gene stability was assessed with geNorm (Qbase+, Biogazelle, Belgium; Vandesompele *et al*. 2002), which calculates the stability of expression (M) for each gene. M*_SDHA_* and M*_RPL13_* were both below 0.7, which is the recommended upper limit for the stability value (M) of a reference gene (Vandesompele *et al*. 2002). The reference gene expression (Cq) did not differ between the treatment groups (ANOVA-test: *SDHA*: F_3,29_=0.31, p=0.81, *RPL13*: F_3,29_=1.12, p=0.36).

#### 2.2.3 Gene expression: Quantitative PCR

The relative quantity of the reverse transcribed target cDNA was assessed using magnetic induction cycler (micPCR, Bio Molecular Systems) and SensiFAST SYBR Lo-ROX Kit (Bioline). For each gene, samples were analyzed in two 48-well qPCR plates. All biological samples were run as technical duplicates on the same plate. Additionally, pooled samples from four great tit samples were run in quadruplicates to serve as calibrator samples for expression normalization. Each plate also included duplicates of sterile H_2_O as no template controls and RNA samples which were not reverse transcribed as controls. For each well, 5 μl of cDNA (1.2 ng/μl) was combined with 6 µl SensiFAST SYBR Lo-Rox Mix, 0.18 µl forward and reverse primers (300nM) and 0.64 µl sterile H_2_O (V_tot_=12 μl) in strip tubes preloaded with mineral oil. Quantitative PCR was run in the magnetic induction cycler with the following program: 95°C 120s, (95°C 5s, 60°C 20s) x 45.

#### 2.2.4 Gene expression: Gene expression normalization and quality filtering

Each plate was confirmed to have a single amplification peak for each primer set and NT-controls were confirmed to show no amplification. Relative expression for each sample was assessed with Pfaffl method (Pfaffl 2001) using the formula below:

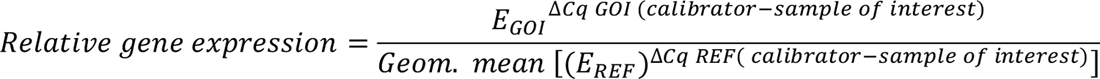

E refers to the average efficiency for each gene in each plate (theoretical maximum would be perfect doubling at each PCR cycle = 2). Efficiencies were obtained from micPCR Software output, which calculates them using the LinRegPCR algorithm described by Ramakers *et al*. (2003) with the formula: *E* = 10^*slope*^ – 1. The slope is determined with linear regression from the amplification curve. Cq is the quantification cycle value for each sample as the number of cycles needed for the fluorescence (describing PCR product quantity) to reach a threshold set by the LinRegPCR algorithm (Ruijter *et al*. 2009). The calibrator Cq is the pooled sample in each run.

Relative gene expression for each individual was calculated as the mean relative expression for the technical duplicates, which was log_2_ transformed for further statistical analyses. Samples with over 30% CV between the relative expression values of the technical duplicates were discarded. Model residuals showed no outlier samples.

### 2.3 Statistical analysis

All statistical analyses were conducted in R Studio (version 4.0.3, R Core Team 2020). To examine variation in breath rate, DNA methylation and gene expression, we used linear (mixed) models, using base R and package *lme4* (Bates *et al*. 2015), while type III ANOVA were calculated using the package *lmerTest* (Kuznetsova et al. 2017) and *car* (Fox & Weisberg 2019). We inspected the normality and homogeneity of variance visually from the model residuals. F-statistics (with associated degrees of freedom) and p-values from type III ANOVA were calculated with the Kenward-Roger method for the mixed models (breath rate and methylation analysis). Random effect significance was calculated using likelihood ratio test by comparing models with and without the random effects. Post-hoc comparisons were assessed with package *emmeans* (Lenth 2021) using Tukey’s multiple comparison procedure. *emmeans* was also used to calculate effect sizes of the hormone treatments. R packages *ggplot2* (Wickham 2016) and *ggpubr* (Kassambara 2020) were used to create figures.

The hormone treatments were considered 2-level factors (CORT yes/no and TH yes/no). CORT, TH and their interaction CORT*TH were fixed effects in all the models examining the effects of hormone treatment. Model covariates were selected based on biological hypotheses. Non-significant interactions were removed to avoid overfitting in all of the models.

For modeling breath rate at 14 days post-hatch, the model included the following covariates that were hypothesized to explain variation in individual metabolism and stress response: wing length (proxy of individual structural size), brood size at 2 days post-hatch (proxy of parental condition and nestling environment) and hatching date (proxy of parental condition and food availability). Breath rate -models included nest ID as a random effect to account for non-independence between individuals from the same nest. As 37 out of the 83 individuals in the breath rate analyses were not molecularly sexed, and sex did not have a significant effect on breath rates in this subset of sexed individuals (F_1,29.7_=0.47, p=0.50), we did not include sex in this model.

For modeling the longitudinal DNA methylation measurements, the fixed effects, in addition to the hormone treatments, were sex, age (categorical variable: 7 or 14 days after hatching, or juvenile), the interaction between age and treatment (since the hormonal treatment may have distinct effects at different developmental stages), CpG-site identity, and the interaction between CpG site and hormonal treatment (since methylation at different genomic positions may have different consequences on gene expression, projecting into the phenotype). Sex was also included in the model as a fixed effect since the target genes were hypothesized to have sex-specific expressions (Nätt *et al*. 2014) and all the individuals were sexed. Due to a relatively small sample size (96 samples from 39 individuals) and many levels of CpG-site identity consuming the degrees of freedom, we did not add other covariates or non-significant interactions to the model at the expense of overfitting. Random effects in the DNA methylation model were sample ID (12 and 4 CpG-sites from the same sample for *NR3C1* and *THRB*, respectively), and individual ID (2 or 3 longitudinal samples from the same individual).

For modeling gene expression measured only at 14 days post-hatch, the response variable was the log_2_ transformed relative gene expression. We included sex (as all individuals were sexed) as a fixed effect in the model in addition to the hormone treatments. As for the breath rate analysis, we included wing length, brood size at day 2 and hatching date as covariates in the model. The gene expression analysis did not include random effects since there were no repeated measures as only one individual at a single time point was included in this analysis as well as methylation analysis.

Furthermore, we hypothesized some associations between breath rates, methylation and gene expression. Gene promoter methylation was hypothesized to correlate negatively with gene expression (Bird 2002). Additionally, breath rate was assumed to correlate with receptor gene expression (reviewed in Kapoor et al. 2006). For both genes, the effect of mean methylation per sample on their respective log_2_ transformed relative gene expression, and the effect of log_2_ transformed relative gene expression on breath rates was examined with Pearson’s correlation at 14 days post-hatch.

## 3. Results

### 3.1. Breath rate

Prenatal corticosterone treatment significantly increased breath rate during handling 14 days after hatching (Figure 1A, Table 5; effect size = 0.55). Neither prenatal thyroid hormone nor the interaction between prenatal corticosterone and thyroid hormones had a significant effect on breath rates (Table 5). While brood size and hatching date were not significantly associated with breath rate, individuals with longer wings (a proxy of individual size) had a marginally higher breath rate (Table 5; estimate**±**SE=1.60**±**0.89, although non-significant). The nest identity accounted for 11.6% of the variance in breath rate, which was not significant (Table 5, Table S2).

**Figure 1.**
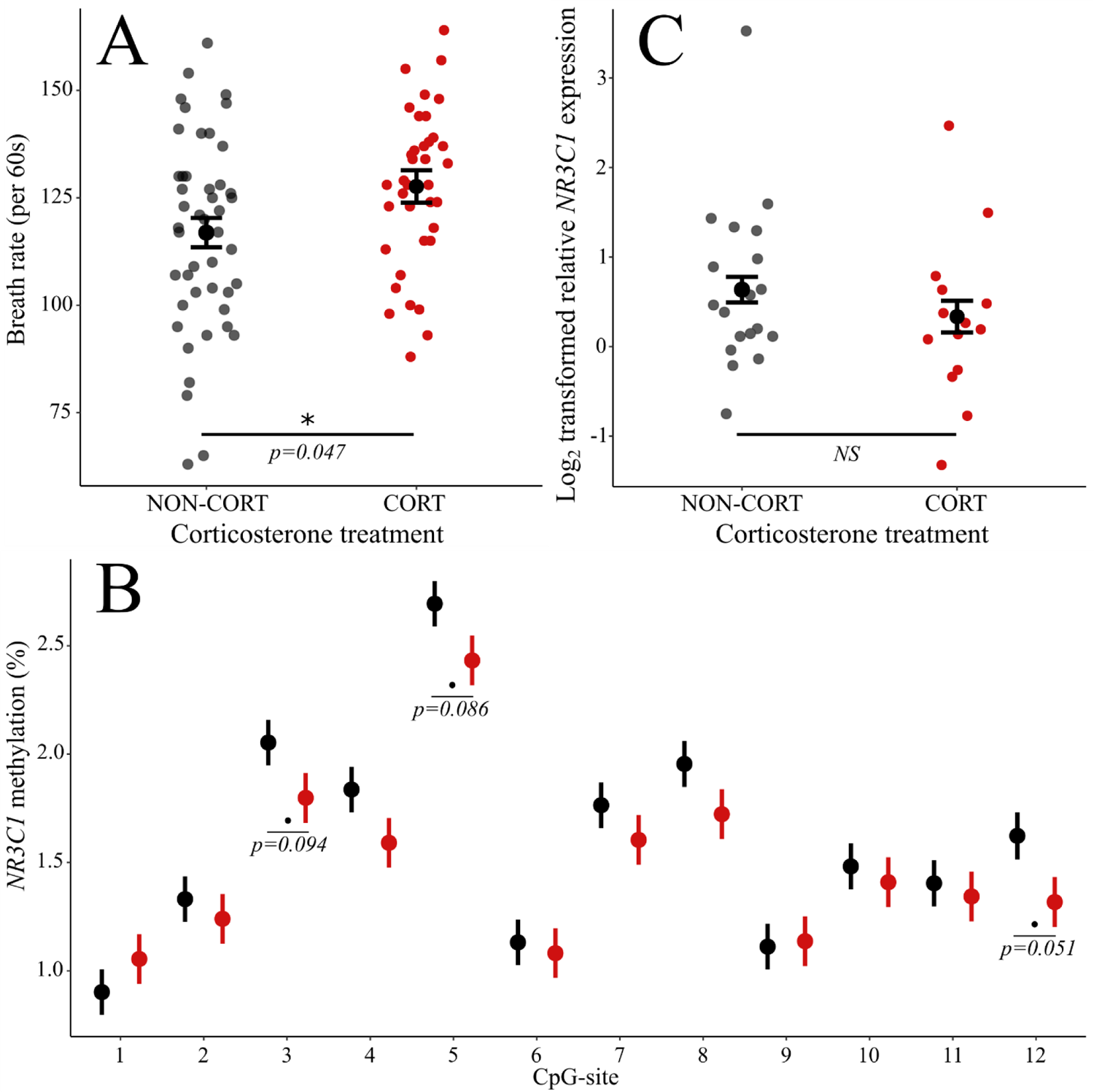
Effects of prenatal corticosterone manipulation on (A): breath rate 14 days after hatching (per 60s), (B): DNA methylation (%) at the quantified 12 CpG-sites of *NR3C1* promoter region (black=NON-CORT; red=CORT) since a significant CORT*Site interaction was detected, and (C): glucocorticoid receptor *NR3C1* relative gene expression in blood cells,. Estimated marginal means and standard errors are given (A-C), with the raw data (A-B). P-values for the effect of corticosterone from type III ANOVA (A-B) and site-specific post-hoc comparison with Tukey’s test (C) are also shown for p<0.10.

**Table 5.**
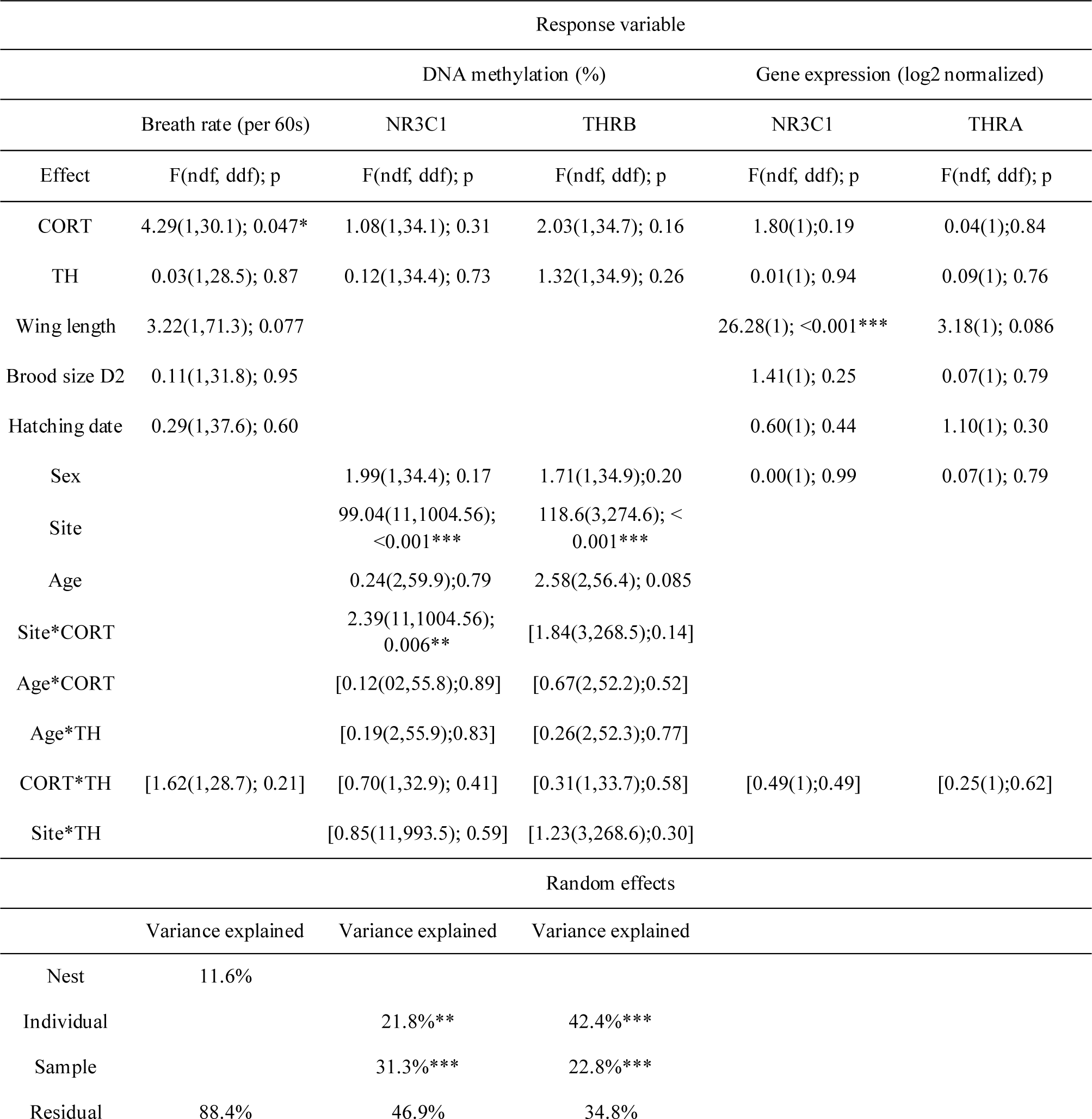
General linear (mixed) model explaining variation in breath rate, DNA methylation, and gene expression. For fixed effects, Type III ANOVA F-statistics, associated degrees of freedom, and p-values are presented. Mixed models (breath rate and DNA methylation) are fit by REML, and degrees of freedom are estimated with Kenward-Roger’s method. For random effects, the percentage of variation explained (VE), and a test of significance (likelihood ratio test, with χ 2 (df) and p-value) are provided (see supplementary Table S2). Significant effects are marked with asterisks (*: p<0.05; **:p<0.01;***:p<0.001). Brackets [] indicate non-significant interaction terms removed from the final model. NOTE: The test statistics for the main effects were from the final models without interaction terms. The exception was the model for NR3C1 DNA methylation, which contains a significant interaction effect between corticosterone (CORT) and CpG-site. Thus, the test statistics for the main effect of CORT represent the contrast with NON-CORT at CpG-site 1, and the main effect of CpG site represents the CpG-site difference in the non-CORT group, respectively. See section 3.2 and Figure 1B for post-hoc comparisons.

### 3.2. DNA methylation

Distinct CpG-sites differed in their methylation value, with site explaining significant variation in DNA methylation for both *NR3C1* and *THRB* (Table 5, Figure 1B). Prenatal corticosterone treatment had CpG-site-specific effects on *NR3C1* promoter methylation (Figure 1B, Table 5; significant CORT*CpG-site interaction). Tukey post-hoc comparisons revealed that for CpG-sites 3, 5 and 12 of the target region, corticosterone decreased DNA methylation coming close to significance (all p<0.094; Figure 1B; effect size= CpG-site 3: −0.59, CpG-site 5: −0.61, CpG-site 12: −0.71). Thyroid hormone treatment, age (Figure 2A), sex, or the interactions between CORT and TH, age, and hormonal treatment, as well as CpG-site and TH had no significant effect on DNA methylation at the *NR3C1* promoter region. A significant amount of variance (conditional on fixed effects) was explained by sample identity (12 CpG-sites from the same sample, 31.3%) and individual identity (2 or 3 longitudinal samples from the same individual, 21.8%) (Table 5).

**Figure 2.**
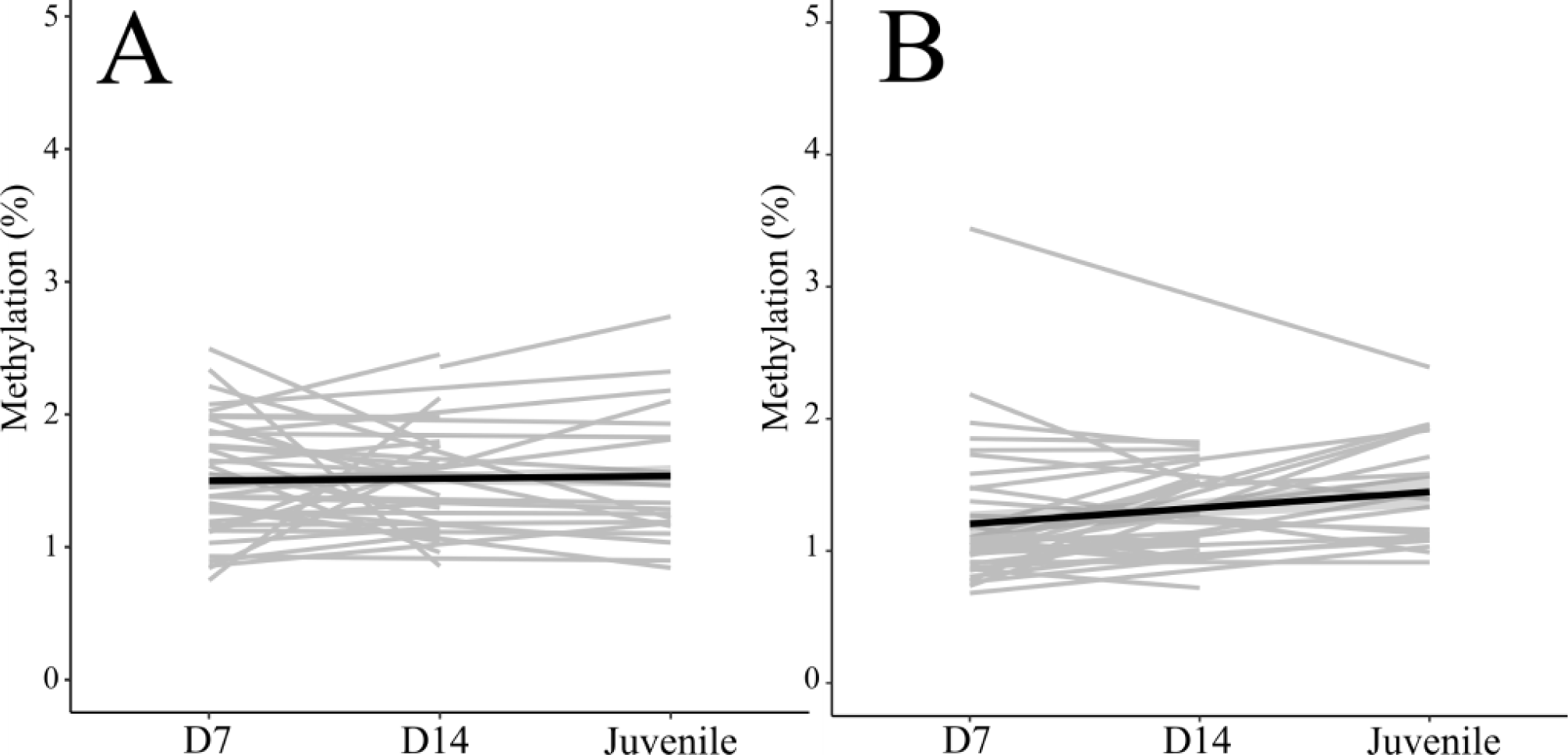
2A: Average methylation percentages pooled across different treatment groups at different ages for glucocorticoid receptor gene *NR3C1*. 2B: Average methylation percentages pooled across different treatment groups at different ages for thyroid hormone receptor gene *THRΒ*. Black lines represent changes in methylation percentage in the overall mean (across all samples), and the grey lines represent changes in methylation percentage (averaged over all CpG sites) for each individual across ages.

For *THRB*, DNA methylation tended to vary with age (p<0.10; Table 5; Figure 2B; estimate**±**SE=Age D7: −0.24**±**0.11, Age D14: −0.17**±**0.11 compared to juveniles). Thyroid hormone treatment, sex, or the interactions between CORT and TH, age and hormonal treatment, CORT and CpG-site or TH and CpG-site did not significantly affect *THRB* methylation (Table 5). Individual identity and sample identity explained a significant amount of variance in *THRB* methylation, 22.8% and 42.4%, respectively (Table S2).

### 3.3. Gene expression

Neither corticosterone (Figure 1C, effect size= *NR3C1*: −0.50, *THRA*: 0.08) nor thyroid hormone treatment had a significant effect on gene expression 14 days post-hatch (Table 5). For *NR3C1,* wing length was significantly negatively associated with gene expression (Table 5, estimate **±**SE= −0.14**±**0.028), whereas sex, brood size and hatching date had no significant effect on gene expression.

For *THRA*, only wing length tended to have a significant negative relationship with gene expression (estimate**±**SE=-0.093**±**0.052, although not significant), whereas no other variable exhibited a significant relationship with *THRA* gene expression (Table 5).

### 3.4. Correlations

While analyzing possible relationships between the variables of interest at day 14 post-hatch, we found *NR3C1* gene expression to have a significant negative bivariate correlation with breath rates (Figure 3A; R(Pearson)=-0.58, p=0.022). Furthermore, *NR3C1* sample mean methylation had a positive bivariate correlation with gene expression (Figure 3B; R(Pearson)=0.46; p=0.007). *THRA* expression showed no significant relationships with breath rates or *THRB* methylation.

**Figure 3.**
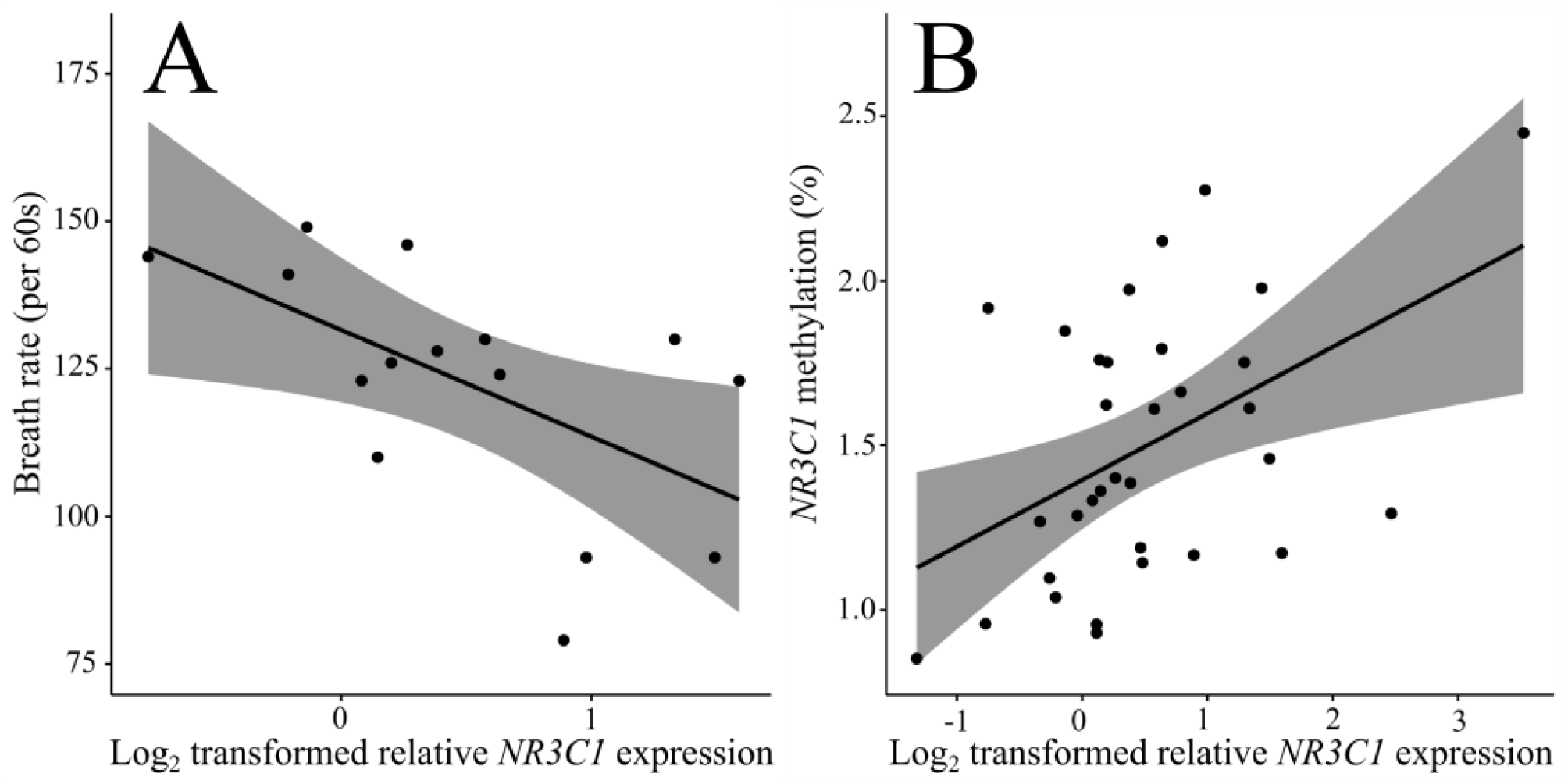
3A: Correlation between breath rate (per 60s) and *NR3C1* gene expression. 3B: Correlation between *NR3C1* promoter methylation (%) and relative gene expression. Regression line and 95% confidence limits are given with the raw data.

## 4. Discussion

Elevation of prenatal corticosterone increased breath rates in great tits 14 days after hatching. Prenatal corticosterone also tended to decrease GCR gene *NR3C1* promoter methylation in 3 of the 12 studied CpG sites, but no age-specific patterns were observed. Prenatal corticosterone had no significant effect on gene expression of GCR (although effect size was of similar magnitude as for breath rate or methylation) or THR, while GCR gene expression was negatively associated with breath rates and showed a strong association with both wing length and hatching date. Elevation of prenatal THs did not influence breath rate, THR methylation or gene expression.

### 4.1 The effects of prenatal hormones on breath rates, DNA methylation and gene expression

#### 4.1.1 Prenatal corticosterone treatment

In line with our hypothesis and previous studies (e.g. Tilgar et al. 2016, *reviewed in* Thayer et al. 2018), prenatal supplementation of great tit eggs with corticosterone significantly increased breath rate, a measure of stress response and metabolism, at 14 days after hatching. Increased breath rate may result from enhanced glucocorticoid response (Carere & can Oers 2004), which was not directly evaluated in this study, yet the evidence is equivocal: in ovo corticosterone treatment has been found to both increase (Freire et al. 2006; Haussmann et al. 2012; Marasco et al. 2012; Ahmed et al. 2014) and decrease (Hayward et al. 2006; Love & Williams 2008; Tilgar et al. 2016) HPA activity and baseline as well as stress-induced corticosterone levels. Furthermore, Podmokla et al. (2018) concluded in a meta-analysis that experimental corticosterone treatment yields neither overall nor manipulation-specific effects on offspring traits. The discrepancy in previous studies show that HPA axis regulation may be subject to maternal corticosterone, affecting stress response and metabolism, yet there may be other biological mechanisms involved and the effects are likely timing-, context- and dose-dependent. The current study elucidates this discrepancy showing that in altricial species such as great tit, pre-incubation corticosterone can increase offspring breath rates and thus, may alter offspring HPA axis reactivity, highlighting the role of maternal effects in shaping offspring phenotype.

The molecular mechanism underlying the observed hormonal effects on breath rates may be related to epigenetic changes since our corticosterone treatment also had CpG-site specific effects, mainly decreasing DNA methylation in the putative promoter area of the glucocorticoid receptor (GCR, *NR3C1*) gene. In line with our results, Ruiz-Raya et al. (2023) found prenatal exposure to alarm calls to reduce GCR promoter methylation in yellow-legged gulls (*Larus michahellis*). However, these results on the site-specific decreased methylation at GCR gene promoter after corticosterone treatment are not in agreement with our primary hypothesis nor with the few previous studies investigating prenatal and early life stress, and GCR methylation: In domestic chickens, a high concentration of corticosterone injected into eggs around mid-incubation increased hypothalamic GCR methylation (Ahmed et al. 2014). Bockmühl et al. (2015) found that postnatal early life stress in mice increased hypothalamic CpG-island shore methylation at certain CpG-sites at GCR. Jimeno et al. (2019) observed an increase in blood GCR promoter methylation resulting from postnatal early life adversity. Azar et al (2022) reviewed prenatal maternal stress to increase offspring peripheral DNA methylation of the GCR gene. Yet, to our knowledge, this is the first study assessing the effects of pre-incubation corticosterone injection on the methylation status of this gene in blood.

Corticosterone treatment did not influence GCR expression significantly, though the effect size was of the same magnitude as for the effects of corticosterone treatment on breath rates and GCR methylation (∼0.5)). Previous studies have found prenatal stress to alter GCR expression, yet the direction of the change has not been unequivocal (Kapoor et al. 2006; Cottrell & Seckl 2009; Zimmer et al. 2017; Ruiz-Raya et al. 2023). As GCR methylation and gene expression were positively correlated in our study, it could be that prenatal corticosterone causes site-specific GCR methylation alterations that would have consequences on gene expression. This complex, possibly activating role of methylation at some CpG-sites has some support from previous literature: Bockmühl et al. (2015) found early-life stress to increase GCR expression by site-specific CpG island shore hypermethylation. In yellow-legged gulls, Ruiz-Raya et al. (2023) found no correlation between GCR expression and average promoter methylation levels, or CpG-site-specific promoter methylation, yet they did find GCR expression to associate with principal component 2 derived from methylation data, which further supports the multifaceted role of promoter methylation in transcriptional regulation. Yet, the positive relationship between methylation at regulatory CpGs and gene expression contrasts the canonical view of the suppressive role of promoter methylation (Bird 2002) and findings of the few previous studies assessing perinatal stress, GCR methylation and expression (Ahmed et al. 2014; Jimeno et al. 2019).

Taken together, increased maternal corticosterone may increase offspring stress response. As DNA methylation was decreased after corticosterone treatment, and changes in the promoter region of GCR did not directly translate to significant differences in GCR expression, the molecular mechanisms remain to be fully elucidated. Our results are in line with those from the same field experiment investigating effects of prenatal corticosterone and thyroid hormones on mitochondrial aerobic metabolism, growth and survival (Cossin-Sevrin et al. 2022). Prenatal corticosterone treatment led to decreased mitochondrial metabolism: It is possible that prenatal corticosterone treatment altered mitochondrial metabolism through glucocorticoid signaling in our study, since glucocorticoid signaling is suggested to alter mitochondrial traits (Casagrande et al. 2020; Ridout et al. 2020).

#### 4.1.2 Prenatal thyroid hormone treatment

Prenatal TH supplementation of great tit eggs did not significantly alter breath rates, GCR or THRB DNA methylation status, or gene expression at the GCR or THRA genes. There are several plausible, mutually non-exclusive, explanations for this. First, it could be that prenatal THs do not have a strong effect on offspring hormonal signaling and stress-related phenotype. The lack of effects of prenatal TH supplementation are in line with Cossin-Sevrin et al. (2022), who also did not find a significant effect of thyroid hormones on growth or mitochondrial metabolism (however, they did find an effect on developmental time), suggesting that experimental corticosterone may have stronger leverage on offspring stress-related phenotype than TH with this experimental set-up. Second, the genes we analysed might not be targets of prenatal THs and their actions on other biological pathways (Vitousek et al. 2019). Third, the effects of maternal hormones are dependent on the expression of transport molecules, cell membrane transporters and deiodinases facilitating the conversion of TH between the inactive and active forms (McNabb & Wilson 1997; Ruuskanen & Hsu 2018). In chickens, a high level of expression of deiodinase (DIO) type 3 by the yolk sac membrane was found since embryonic day 5 (Too et al. 2017), which might have some function in de-activating excessive THs. In passerines, Ruuskanen et al. (2022) also found early-stage embryos to express DIO2, DIO3, THRA, THRB and monocarboxyl membrane transporter MCT8, suggesting that altricial embryos are also able to modulate the effects of egg TH during embryonic development. Fourth, prenatal TH elevation may have tissue-specific effects on gene methylation and expression, such as brain, but not in blood cells (McCormick et al 2000; Bockmühl et al. 2015; Lattin et al. 2015; but see support for between-tissues correlations in Daskalakis et al 2014). Yet, even if blood and brain hormone receptors are not tightly correlated, information on blood levels may provide valuable functional information (Jimeno & Zimmer 2022). Fifth, prenatal THs work in synergy with hormones that were not included in this study, for example, Wang et al. (2007) found that oral dosing of thyroid hormone (T3) together with growth hormone injections had synergist effects on body fat and hepatic gene expression of juvenile chickens.

### 4.2 Patterns of DNA methylation

The overall methylation percentages for both genes, glucocorticoid receptor and thyroid hormone receptor, were generally low (median < 2%). These results corroborate with previous findings from birds where CpG-dense promoters and transcription start sites are less methylated specifically for the gene coding for GCR (yellow-legged gulls: Ruiz-Raya et al. 2023), as well as for genome-wide patterns (great tits: Derks et al. 2016; Laine et al. 2016). Derks et al. (2016) found that great tit transcription start sites in the brain and blood are generally lowly methylated in the tissues in which they are anticipated to be expressed. Both genes of interest, glucocorticoid receptor GCR and thyroid hormone receptor THRΒ, exhibited statistically significant differences between the methylation percentages of individual CpG-sites. These results suggest that certain CpG-site methylation may be more important in the regulation of gene expression rather than the average methylation percentage of a certain CpG-island. Indeed, it has been shown that even within promoters, the entire sequence may not be methylated the same way and short sequences may have distinct methylation patterns depending on which transcription factors bind which sites (Tohgi et al. 1999).

For GCR, a large proportion of variance in methylation percentages, conditional on fixed effects, was explained by sample identity (i.e., 31.3%), implying consistency in the methylation percentage between different CpG sites of the same sample, probably partly due to co-methylation over neighboring CpG-sites (Eckhardt et al. 2006; Jimeno et al. 2019). This may be due to sample-specific blood cell type composition, which varies between individuals, although the vast majority of avian blood DNA is from nucleated red blood cells (Husby 2020). A significant amount of variance (i.e. 22.0%) was also explained by individual identity, which demonstrates consistent inter-individual differences in methylation through time. For THRΒ, the largest proportion of variance was explained by individual identity (42.4%), but sample identity also explained a substantial part of the variance (22.8 %). The relatively high within-individual consistency in methylation percentages suggested that methylation patterns for different individuals had persisted from 7 days of age to juvenility. This indicates relatively robust and consistent methylation in the analyzed regions across time. In contrast, Bockmühl et al. (2015) found an age-related (10-days-vs. 6- weeks- vs. 3-months-old) increase in methylation in certain CpG-sites in the shore region of a CpG-island upstream from GCR in the hypothalamus of mice that had been exposed to early life stress. Early-life stress induced an increase in the overall methylation of this CpG-island shore was only observed at the later age (Bockmühl et al. 2015). In turn, Marasco et al. (2012) injected corticosterone to the eggs of Japanese quail and found a hyper-regulated HPA response as elevated circulating corticosterone levels during acute stress at 64 days after hatching, but not at 22 days after hatching. The results of these previous studies suggest that the effects of prenatal hormonal treatments might be more evident at adulthood than during postnatal development. Consequently, the time interval between prenatal hormonal injection and post-hatching days 7 and 14 was maybe too short to detect possible differences in methylation percentages resulting from prenatal hormonal treatment. However, we found no significant impact in juveniles (approximately 4-months-old) either, but the number of juveniles for each treatment group was only 5 (4 for THRΒ CORT group), which may have been a sample size too small to detect treatment differences in methylation percentages.

### 4.3 Patterns of gene expression

There was a negative relationship between body size (i.e. wing length) and GCR gene expression, larger individuals having lower GCR expression levels. Nutritional or developmental stages may explain the relationship between body size and GCR expression. Food insecurity and malnutrition has been found to stimulate stress and increase cortisol levels in humans (Sawaya 2006; Freitas et al. 2018), which in turn could influence stress hormone receptor expression (Cottrell & Seckl 2009). Alternatively, it may be that stress responsiveness, as altered GC-receptor expression, may influence energy resources allocated between stress and growth, seen as varying growth rates. Thus, differences in gene expression of the GC-receptor may drive differences in growth rates.

In contrast to GCR expression, THRΑ expression did not correlate with any of the studied biological and ecological covariates (wing length, hatching date, brood size, breath rate). A variety of factors could alter thyroid hormone levels of the nestling’s mother’s plasma, such as food and iodine availability, endocrine-disrupting molecules, stress, and therefore indirectly also pathogens and intra- and interspecies interactions (Ruuskanen & Hsu 2018). Therefore, a robust regulation in TH receptor expression may be adaptive to protect the developing individuals from environmental and/or physiological variation both pre- and post-hatch.

## 5. Conclusions

This study supports the view that maternal corticosterone may influence offspring metabolism and stress response via epigenetic alterations, yet their possible adaptive role needs to be further tested by assessing long-term fitness consequences of these effects across different environments. Further, we encourage future research to analyze whole-genome methylation patterns and transcriptomic profiles to elucidate the pathways linking prenatal hormonal exposure and postnatal HPA response.

## Acknowledgements

We are grateful to Meri Lindqvist for guidance with bisulfite conversion and Jorma Nurmi for help in the fieldwork.

The study was funded by KLYY foundation (MH), Valto Takala foundation (MH), FIMM-EMBL rotation program (MH), Jenny and Antti Wihuri foundation (MK), Maupertuis Grant (NCS), University of Turku Graduate School (NCS), Ella and Georg Ehrnrooth Foundation (BYH), Turku Collegium for Science and Medicine (AS), European Commission Marie Sklodowska-Curie Fellowship #894963 (AS) and Academy of Finland (#286278 to SR, #332716 to BYH and #325937 to RK).

## 6. Author contribution statement

SR, AS, BYH conceived the study. All authors participated in data collection (field work and/or laboratory methodology). MH, SR, AS, BYH conducted the statistical analyses. MH wrote the original draft, which all authors edited.

## 7. Data availability statement

All data, and R scripts are archived and available in Figshare (DOI: 10.6084/m9.figshare.22153001).

## 8. Supplementary material

Supplementary materials include Tables S1 and S2 (submitted as a separate file).

## 9. Ethics

The authors confirm that the manuscript has not been submitted elsewhere and that all research meets the ethical guidelines of the study country, Finland. All procedures were approved by the Animal Experiment Committee of the State Provincial Office of Southern Finland (license no. ESAVI/5718/2019) and by the Environmental Center of Southwestern Finland (license no. VARELY/924/2019) granted to SR.

[Table S1 legend] Table S1. Sex ratios: number of individuals (F:M) included in the analysis after quality filtering for each gene (NR3C1=glucocorticoid receptor; THRB=thyroid hormone receptor β), treatment group (CO=control; CORT=corticosterone; CORT + TH=corticosterone and thyroid hormone; TH=thyroid hormone) and age (DAH=days after hatching).

[Table S2 legend] Table S2. Estimated proportion of variance (%Var) explained by random effects for all models. The proportion of explained variance was calculated as the estimated variance for the random factor in question divided by the total variance. Significance tests were perfomed by log-likelihood ratio test between models with different random structures. Significance is marked with asterisks (*: p<0.05; **:p<0.01;***:p<0.001).

## Notes

https://doi.org/10.6084/m9.figshare.22153001.v1

### Competing Interest Statement

The authors have declared no competing interest.

