## Supplementary Materials for "From maternal glucocorticoid and thyroid hormones to epigenetic regulation of gene expression: an experimental study in a wild bird species"

Table S1. Sex ratios: number of individuals (F:M) included in the analysis after quality filtering for each gene (*NR3C1*=glucocorticoid receptor; *THRB*=thyroid hormone receptor β), treatment group (CO=control; CORT=corticosterone; CORT + TH=corticosterone and thyroid hormone; TH=thyroid hormone) and age (DAH=days after hatching).

|  | NR3C1 | | | | |
| --- | --- | --- | --- | --- | --- |
| Age | CO | CORT | CORT + TH | TH | Total |
| 7 DAH | 10 (4:6) | 9 (2:7) | 9 (1:8) | 10 (4:6) | 38 (11:27) |
| 14 DAH | 10 (4:6) | 10 (3:7) | 8 (1:7) | 10 (4:6) | 38 (12:26) |
| Juvenile | 5 (2:3) | 5 (2:3) | 5 (1:4) | 5 (1:4) | 20 (6:14) |
| Total | 25 (10:15) | 24 (7:17) | 22 (3:19) | 25 (9:16) | 96 (29:67:2) |
|  | THRB | | | | |
| Age | CO | CORT | CORT + TH | TH | Total |
| 7 DAH | 10 (4:6) | 10 (3:7) | 8 (1:7) | 10 (4:6) | 39 (12:26) |
| 14 DAH | 10 (4:6) | 10 (3:7) | 7 (0:7) | 10 (4:6) | 38 (11:26) |
| Juvenile | 5 (2:3) | 4 (1:3) | 5 (1:4) | 5 (1:4) | 19 (5:14) |
| Total | 25 (10:15) | 24 (7:17) | 22 (2:18) | 25 (9:16) | 94 (28:66) |

Table S2. Estimated proportion of variance (%Var) explained by random effects for all models. The proportion of explained variance was calculated as the estimated variance for the random factor in question divided by the total variance. Significance tests were perfomed by log-likelihood ratio test between models with different random structures. Significance is marked with asterisks (*: p<0.05; **:p<0.01;***:p<0.001).

|  | Breath rate | | | | | | | | | |
| --- | --- | --- | --- | --- | --- | --- | --- | --- | --- | --- |
| Random effect(s) | %Var | Model | Parameters | AIC | Log likelihood | Deviance | Test | χ2 | Δdf | p |
| None |  | 1 | 7 | 745.99 | -366.00 | 731.99 |  |  |  |  |
| Nest | 11.6 | 2 | 8 | 747.76 | -365.88 | 731.76 | 1 vs 2 | 0.2284 | 1 | 0.63 |
| Residual | 88.4 |  |  |  |  |  |  |  |  |  |
|  | DNA methylation: NR3C1 | | | | | | | | | |
| Random effect(s) | %Var | Model | Parameters | AIC | Log likelihood | Deviance | Test | χ2 | Δdf | p |
| Sample | 21.8 | 1 | 30 | 1576.4 | -758.22 | 1516.4 | 1 vs 3 | 243.64 | 1 | <0.001 *** |
| Individual | 31.3 | 2 | 30 | 1811.0 | -875.51 | 1751.0 | 2 vs 3 | 9.052 | 1 | 0.003** |
| Individual and sample |  | 3 | 31 | 1569.4 | -753.69 | 1507.4 |  |  |  |  |
| Residual | 46.9 |  |  |  |  |  |  |  |  |  |
|  | DNA methylation: THRB | | | | | | | | | |
| Random effect(s) | %Var | Model | Parameters | AIC | Log likelihood | Deviance | Test | χ2 | Δdf | p |
| Sample | 22.8 | 1 | 11 | 537.43 | -257.71 | 515.43 | 1 vs 3 | 27.18 | 1 | <0.001*** |
| Individual | 42.4 | 2 | 11 | 557.62 | -267.81 | 535.62 | 2 vs 3 | 47.37 | 1 | <0.001*** |
| Individual and sample |  | 3 | 12 | 512.25 | -244.13 | 488.25 |  |  |  |  |
| Residual | 34.8 |  |  |  |  |  |  |  |  |  |
